# Denisovan Ancestry in East Eurasian and Native American Populations

**DOI:** 10.1101/017475

**Authors:** Pengfei Qin, Mark Stoneking

**Affiliations:** Department of Evolutionary Genetics, Max Planck Institute for Evolutionary Anthropology, 04103 Leipzig, Germany

**Keywords:** Denisovans, Neanderthals, Archaic admixture, Modern humans

## Abstract

Although initial studies suggested that Denisovan ancestry was found only in modern human populations from island Southeast Asia and Oceania, more recent studies have suggested that Denisovan ancestry may be more widespread. However, the geographic extent of Denisovan ancestry has not been determined, and moreover the relationship between the Denisovan ancestry in Oceania and that elsewhere has not been studied. Here we analyze genome-wide SNP data from 2493 individuals from 221 worldwide populations, and show that there is a widespread signal of a very low level of Denisovan ancestry across Eastern Eurasian and Native American (EE/NA) populations. We also verify a higher level of Denisovan ancestry in Oceania than that in EE/NA; the Denisovan ancestry in Oceania is correlated with the amount of New Guinea ancestry, but not the amount of Australian ancestry, indicating that recent gene flow from New Guinea likely accounts for signals of Denisovan ancestry across Oceania. However, Denisovan ancestry in EE/NA populations is equally correlated with their New Guinea or their Australian ancestry, suggesting a common source for the Denisovan ancestry in EE/NA and Oceanian populations. Our results suggest that Denisovan ancestry in EE/NA is derived either from common ancestry with, or gene flow from, the common ancestor of New Guineans and Australians, indicating a more complex history involving East Eurasians and Oceanians than previously suspected.

## Significance Statement

Archaic hominins genetically interacted with the ancestors of present-day humans, but the scope and magnitude of this interaction has not been fully documented. Our study reveals that ancestry from Denisovans, (an archaic human group identified in the Altai Mountains of southern Siberia), is prevalent in populations across eastern Eurasia and the Americas, rather than only in island populations of Southeast Asia and Oceania as previously reported. We show that the Denisovan ancestry in Eastern Eurasians and Native Americans is probably derived from either common ancestry with, or ancient gene flow from, the common ancestor of present-day New Guineans and Australians, thereby demonstrating a more complex relationship of these human populations than previously suspected.

## Introduction

Following the initial description and analysis of a genome sequence from an archaic human fossil from Denisova Cave in southern Siberia (1), Denisovan admixture was subsequently found to be limited to populations from eastern Indonesia, the Philippines, and Near and Remote Oceania (1–3). This finding was quite surprising, given that the Denisova Cave site is located some 7000 km away from the populations that currently exhibit Denisovan ancestry, and was interpreted as suggesting that Denisovan admixture occurred somewhere in the vicinity of island Southeast Asia (2). However, further studies have indicated that Denisovan ancestry may be more widespread than initially thought (4). In particular, a recent study inferred low levels of Denisovan ancestry of about 0.2% in three genome sequences, from a Dai and a Han Chinese from East Asia and a Karitiana from South America (5). Moreover, it appears that a genetic adaptation to high altitude in the Tibetan Plateau occurred via introgression from a Denisovan-related population into ancestral Tibetans (6). Overall, these results suggest that Denisovan ancestry is not limited to populations from island Southeast Asia and Oceania, as originally thought. However, only a few populations have been systematically evaluated for signals of Denisovan ancestry; it is not clear if a very low level of Denisovan ancestry is geographically widespread, or rather limited to only a few populations outside of island Southeast Asia and Oceania. An additional question of interest is whether the Denisovan ancestry in these other populations reflects the same admixture event that contributed Denisovan ancestry to island Southeast Asian and Oceanian populations, or a different admixture event.

To address these and other questions related to Denisovan ancestry in human populations, we here present a systematic investigation of Denisovan introgression in Eastern Eurasia (defined here to include South Asia, East Asia, Southeast Asia, Central Asia, Siberia, and Oceania) and in Native American populations, hereafter abbreviated as EE/NA. Genome-wide data were collected from worldwide populations and analyzed along with high coverage genomes sequences from a Denisovan (3) and a Neanderthal (5). The analyses we report provide more details concerning the admixture history of modern humans with Denisovans.

## Results

### Relationship of present-day humans and archaic hominins

We assembled a dataset which, after quality filtering, consisted of 2493 individuals from 221 populaions, all genotyped on the Affymetrix Human Origins Array (8). After merging the human data with the chimpanzee, Denisovan, and Neanderthal genome sequences, there were nearly 600,000 SNPs for analysis. To investigate the relationship of the diverse present-day human populations relative to archaic hominins, we carried out Principal Component Analysis (PCA) (9) on the chimpanzee, Neanderthal and Denisovan data, and projected the modern human samples onto the plane defined by the top two eigenvectors. The human samples all appear at the center of the plot (Fig. S1*A*); magnification of the central portion of the plot shows that humans separate into three clusters relative to archaic hominins and chimpanzees: African, Oceanian and other Non-African (Fig. S1*B*). To more clearly visualize the patterns, we plotted the mean of eigenvectors 1 and 2 for each of the 221 modern human populations (Fig. 1*A*). The first eigenvector separates the Africans from non-Africans and shows that the non-Africans are clearly closer to archaic hominins than are the Africans. The second eigenvector suggests closer genetic affinity between Oceanians and Denisovan than between other populations and Denisovan. There is a clear cline of Denisovan-related ancestry in Oceanian with Australians and New Guineans having the most Denisovan ancestry (Fig. S2). The Mamanwa, from the Philippines, are also involved in this cline, which is consistent with previous findings that the Mamanwa are related to Australians and New Guineans, and Denisovan admixture occurred in a common ancestral population of the Mamanwa, Australians, and New Guineans (2).

**Fig. 1.**
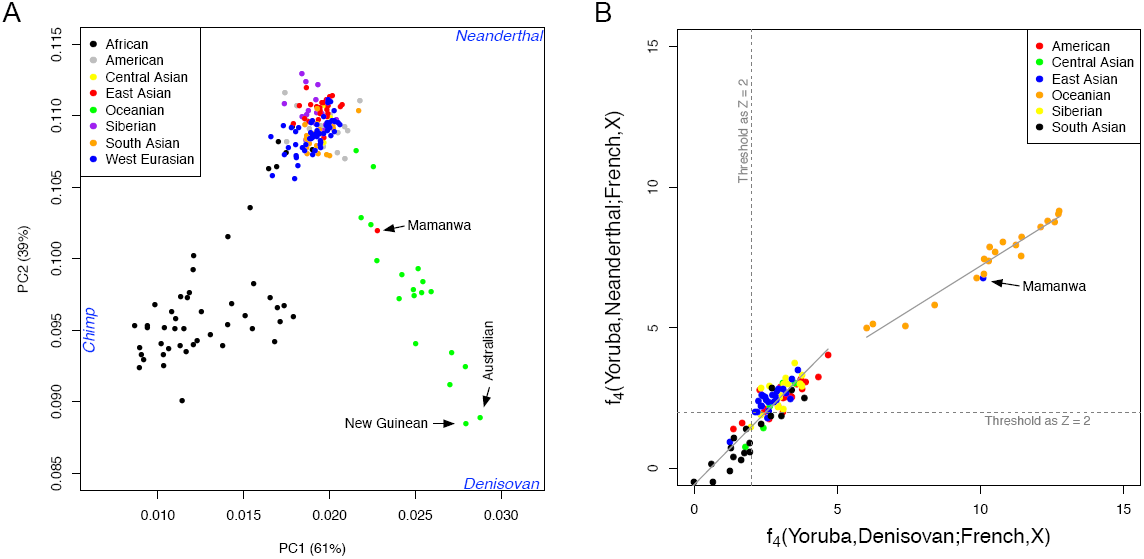
The relationships among modern human populations relative to archaic humans. (A) PCA of 221 populations projected onto the top two eigenvectors defined by Neanderthal, Denisovan and chimpanzee. The mean values of eigenvectors 1 and 2 are plotted for each population. (B) Formal admixture tests suggest a substantial number of EE/NA and Oceanian populations inherited significantly more archaic ancestry than West Eurasian populations. We used *f*_4_ statistics of the form *f*_4_(*Yoruba, Archaic; French, X*) to test admixture between archaic humans and modern human populations. A Z-score larger than 2 was set as the threshold for determining if admixture is significantly greater than zero.

### Additional Archaic Ancestry in EE/NA Populations

PCA is a descriptive analysis that is useful for indicating potential admixture events, but cannot be used to prove that admixture occurred. We therefore applied formal tests to document potential admixture between archaic hominins and modern humans. Since EE/NA populations have on average inherited more archaic ancestry than West Eurasian populations (7), we computed *f*_4_ statistics (8, 10) of the form *f*_4_(*Yoruba, Archaic; French, X*), in which *X* is an EE/NA population. A significantly positive statistic (Z-score> 2) is evidence that EE/NA possesses more Archaic (either Neanderthal or Denisovan) alleles than does the French population. Significantly positive statistics (Z-score> 6) are obtained for all Oceanian populations, which are much higher than those for other EE/NA populations, indicating more archaic ancestry in Oceanians s (Fig. 1*B* and Table S3). Moreover, there are more Denisovan than Neanderthal alleles shared with the Oceanian and Mamanwa populations (Fig. 1*B*), although Neanderthal ancestry is also elevated, probably because signals of Denisovan and Neanderthal ancestry are difficult to distinguish in this analysis. In addition to the Oceanian populations, many additional EE/NA populations exhibit significant Z-scores (> 2), indicating they have more archaic alleles than the French population has. However, unlike the Oceanian populations, the inferred amounts of Denisovan and Neanderthal alleles are approximately the same in these EE/NA populations (Fig. 1*B*). It is thus not clear from this analysis if the additional archaic ancestry in these EE/NA populations reflects Neanderthal ancestry, Denisovan ancestry, or both. In order to increase the power of these tests, we combined the data from the East Asian, Siberian, and native American populations, and obtained significantly higher signals of archaic ancestry (Z-score =3.12 for Neanderthal and 3.64 for Denisovan, compared to the average single population Z-scores of 2.79±0.07 for Neanderthal and 3.17±0.08 for Denisovan). To ensure that our results are not influenced by the choice of African (Yoruba) and European (French) reference populations used in this *f*_4_ analysis, we repeated the analysis with different reference populations and obtained similar results (Fig. S3).

We further computed the statistic *f*_4_(*Yoruba, X; Neanderthal, Denisovan*) which compares the genetic affinity of present-day non-Africans with different archaic hominins (Fig. 2). Positive values of this *f*_4_ statistic indicate excess sharing of Denisovan alleles (relative to Neanderthals); negative values indicate excess sharing of Neanderthal alleles, and values near zero indicate equivalent amounts of alleles shared with Denisovans and Neanderthals. We obtain larger values in Oceanians than in other non-Africans, which is consistent with the observations based on the PCA and on formal tests of admixture. The largest values are observed in New Guineans, Australians, and some populations from Remote Oceania (Table S4), consistent with previous results (2). Negative values are observed in most non-African populations, indicating sharing of more Neanderthal than Denisovan alleles in these populations. Moreover, larger values are observed in East Asians than in West Eurasians, suggesting most East Asians share more alleles with Neanderthal than West Eurasians do. Native Americans are generally more similar to West Eurasians in the patterns of allele sharing, suggesting that either Native Americans have the same amount of allele sharing with Neanderthal as West Eurasians do, or Native Americans shared more Neanderthal alleles than West Eurasians and additionally share a small amount of Denisovan alleles. This procedure was repeated using different African populations in the *f*_4_ statistics and similar results were obtained (Fig. S4).

**Fig. 2.**
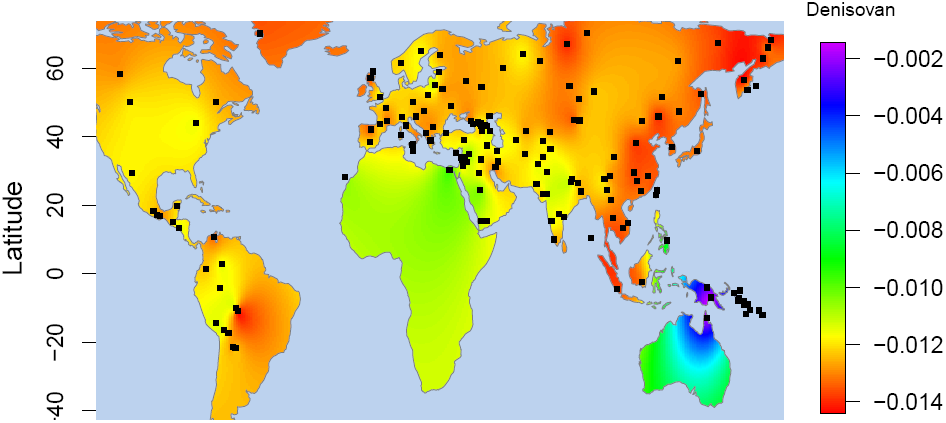
Archaic introgression in modern humans is prevalent and varies across different geographic regions. The sharing of Neanderthal and Denisovan alleles with each non-African population was measured by *f*_4_ statistics of the form *f*_4_(*Yoruba, X; Neanderthal, Denisovan*). An excess of allele sharing with Denisovan yields positive values while an excess with Neanderthal yields negative values. The heat plot values indicated on the map are valid only for regions covered by our samples.

### Denisovan ancestry in Oceanians

Denisovan ancestry in Oceanians has been documented previously (1–3). To verify and extend these previous results, we analyzed a larger set of Oceanian populations that were genotyped on a different platform, and also utilized the high-coverage archaic genomes (3, 5).

Following previous methods (8, 11), we used a ratio of *f*_4_ statistics to estimate the admixture proportion of Denisovans (*р*_*D*_(*X*)) in Oceanians (see *SI*). Since Oceanians retain both Denisovan and Neanderthal ancestry, we used Han Chinese to control for the Neanderthal ancestry in Oceanians, under the assumption that Han and Oceanians share a similar number of Neanderthal alleles. To evaluate the validity of this assumption, we examined the *f*_4_(*Yoruba, Han; Neanderthal, X*) and *f*_4_(*Yoruba, Han; Denisovan, X*) statistics for each Oceanian population *X*. If Han and the *X* possess similar amounts of Neanderthal alleles, then any changes in the two statistics will be driven by the varying amount of their Denisovan alleles. The two statistics will thus have a linear relationship with an intercept close to (0,0). However, if the amount of Neanderthal alleles differs in the two populations, then the linear model will not cross at the point of origin. Empirically, these two sets of *f*_4_ statistics are indeed linearly correlated (*R*^2^ = 0.99; Fig. S5) and the intercept for the linear model fitting the data is near (0,0), indicating that Han and Oceanians are similarly close to Neanderthal. (see *SI*).

The highest *р*_*D*_(*X*) value is observed in Australians and New Guineans (0.034±0.002 and 0.034±0.005 respectively) (Fig. 3*A*), which is consistent with previous results (2) and suggests Denisovan introgression into the common ancestor of Australians and New Guineans. We also observed high amounts (>3%) of Denisovan ancestry in Bougainville, as observed previously (2), and in Santa Cruz, a population from Remote Oceania which was not analyzed previously. This latter result is in keeping with previous observations of extraordinarily high frequencies and diversity of mtDNA and Y-chromosome haplogroups of New Guinean origin in Santa Cruz (12, 13), which suggest high amounts of New Guinean ancestry (and thereby Denisovan ancestry) in Santa Cruz. All other Oceanian populations have Denisovan ancestry ranging from 0.9-3%.

**Fig. 3.**
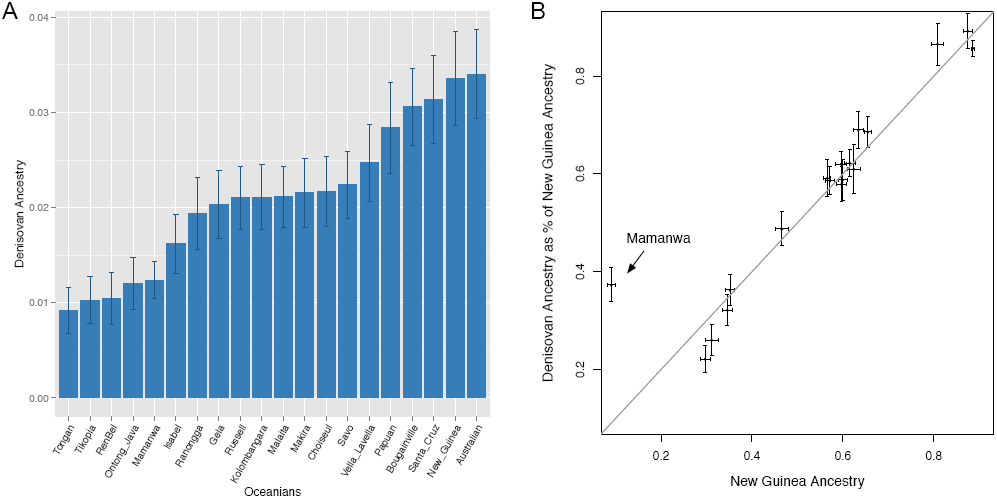
Denisovan ancestry in Oceanians. (A) Estimated Denisovan ancestry in Oceanian populations. (B) Denisovan ancestry in Oceanians is highly correlated with their New Guinean ancestry.

It was shown previously that Denisovan ancestry in Oceanian groups (other than Australia and New Guinea), as well as in eastern Indonesia, was likely to be an indirect consequence of admixture with New Guineans, as the Denisovan ancestry in these other Oceanian and eastern Indonesian groups is proportional to their New Guinea ancestry (2). We observe a similar correlation for the 17 Oceanian populations (excluding Australia and New Guinea) in the present study (Fig. 3*B*). However, given that Australia and New Guinea share common ancestry, it is possible that the Denisovan ancestry in these Oceanian populations was contributed by admixture from Australia or from the ancestral Australia-New Guinea population, rather than directly from New Guinea; these possibilities were not examined in previous studies. We evaluated these alternative possibilities by the statistic *f*_4_(*Yoruba, X; NewGuinea, Australia*), and found that Oceanians share significantly more alleles with New Guineans than with Australians (Table S5). These results suggest that Denisovan ancestry in Oceanians is likely to derive from recent admixture with New Guineans, rather than admixture with Australians or the common ancestor of Australians and New Guineans.

### Denisovan introgression in East/South Asian, Siberian and Native American populations

To detect Denisovan introgression in EE/NA populations, we computed the ratio of two *f*_4_ statistics (*R*_*D*_(*X*)): *f*_4_(*Yoruba, Denisovan, French, X*) and *f*_4_(*Yoruba, Denisovan, French, X*), for each EE/NA population. Populations with *R*_*D*_(*X*) > 1 are likely to have Denisovan ancestry (see *SI*). Before we applied the approach to empirical data, we evaluated the performance of this statistic via simulations (see *SI* and Fig. S6). For simulated populations with Denisovan ancestry, we always observed large ratios (*R*_*D*_(*X*) > 1), while small ratios (*R*_*D*_(*X*) < 1) were obtained for simulated populations without Denisovan ancestry (Fig. S7). We then computed *R*_*D*_(*X*) for all EE/NA populations exhibiting significant admixture signals with Denisovans and/or Neanderthals in formal admixture tests (Fig. 1*B* and Table S3). As expected, we observed large ratios (*R*_*D*_ (*X*) > 1) in all Oceanian populations (Fig. 4). Variable results were obtained for the other EE/NA populations, with *R*_*D*_ (*X*) > 1 observed in several populations, including most Native American populations (Fig. 4). Overall, this analysis indicates that there are EE/NA populations outside Oceania with a clear signal of Denisovan ancestry. Similar results are obtained with the use of different reference populations (Table S6).

**Fig. 4.**
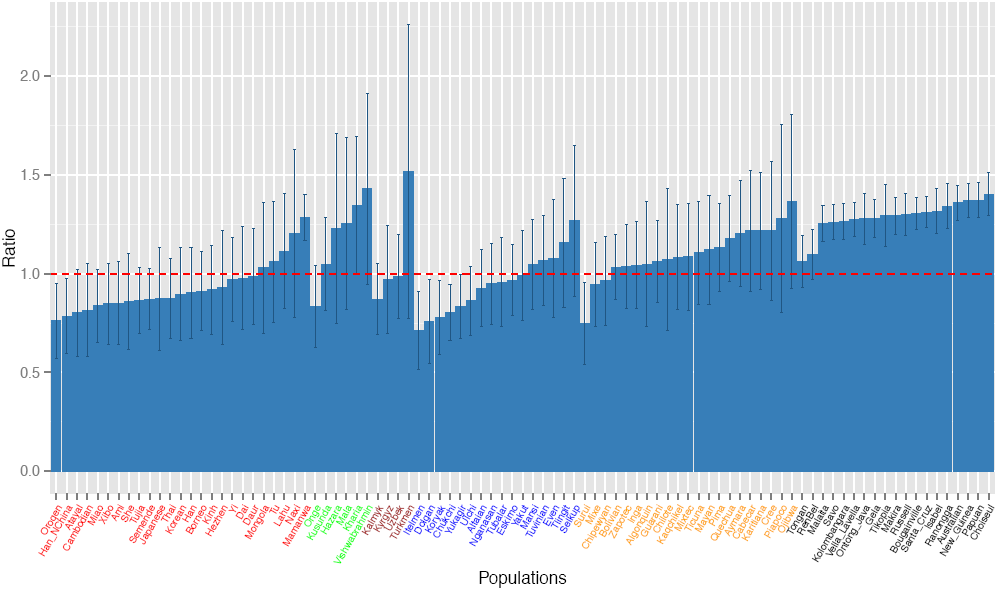
Widespread Denisovan ancestry in EE/NA populations. Values of the *R*_*D*_(*X*) ratio are plotted for all EE/NA populations which give significant signals of admixture with Neanderthal or Denisovan in formal tests; values greater than one (dashed line) are indicative of Denisovan ancestry.

Is this presumptive Denisovan ancestry in EE/NA populations from the same admixture event that contributed Denisovan ancestry to Oceanian populations, or does it rather represent a separate admixture event (or events) between modern humans and Denisovans? We postulated that if it reflects the same event that contributed Denisovan ancestry to Oceanians, then the amount of Denisovan ancestry in EE/NA populations should be correlated with the amount of New Guinean ancestry. Because the estimated amount of Denisovan ancestry is quite small in EE/NA populations, and difficult to distinguish from Neanderthal ancestry, we instead compared the overall amount of archaic admixture in EE/NA populations (as a fraction of that in New Guinean), which is calculated by the ratio of *f*_4_(*Yoruba, Denisovan; French, X*) and *f*_4_(*Yoruba, Denisovan; French, NewGuinean*), to the statistic *f*_4_(*Yoruba, Denisovan; French, X*). These two values are significantly correlated (Fig. 5*A*, Pearson *R*^2^ = 0.23, *p* = 2.5×10^-3^). However, this analysis could be confounded by admixture between East Asians and New Guineans, which has occurred as a consequence of the Austronesian expansion (14, 15) and perhaps other population movements. We therefore removed the East Asian populations and repeated the analysis just for Siberians and Native Americans, and obtained an even higher correlation (Fig. 5*B*, Pearson *R*^2^ = 0.54, *p* = 3.8×10^-6^). Thus, archaic ancestry in EE/NA populations is significantly correlated with their New Guinea ancestry, suggesting that the Denisovan ancestry in EE/NA and Oceanian populations reflects the same admixture event.

**Fig. 5.**
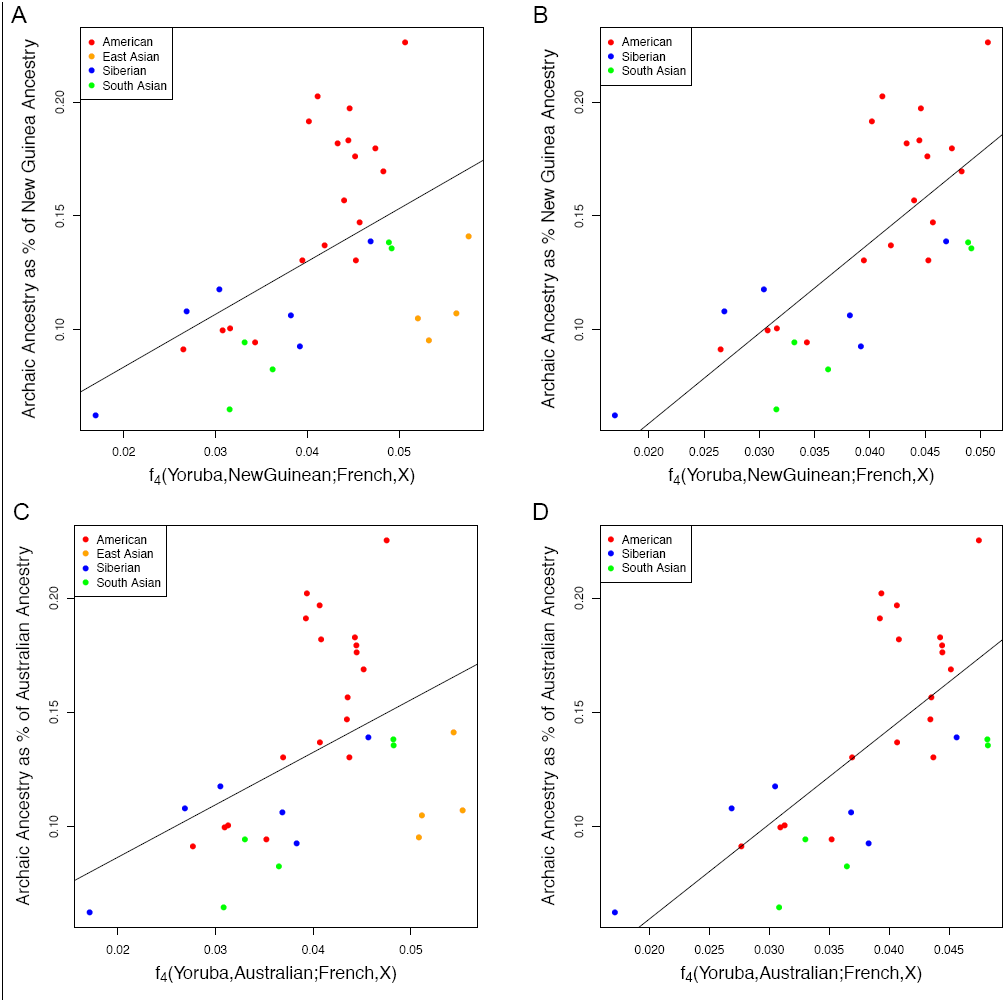
Denisovan introgression in EE/NA populations is correlated with their genetic affinity with Oceanians. Genetic affinity is measured by *f*_4_(*Yoruba, New Guinean; French, X*). We observed a significant correlation between Denisovan ancestry and New Guinea ancestry with (A) *R*^2^= 0.23 (*р* = 2.5×10^-3^) for all EE/NA populations with *R*_*D*_(*X*) > 1, and (B) *R*^2^ = 0.54 (*р* = 3.8×10^-6^) when we remove East Asians. Significant correlations were also observed when New Guinean was replaced by Australian with (C) *R*^2^= 0.19 (*р* = 6.7×10^-3^) and (D) *R*^2^= 0.48 (*р* = 2.0×10^-5^) when we remove East Asians.

However, there are (at least) two potential alternate scenarios that could explain these results. First, Denisovan admixture could have occurred in a population that was ancestral to both EE/NA and Oceanian populations; second, admixture could have occurred in a population that was ancestral specifically to Mamanwa, Australians, and New Guineans (as suggested previously (2)), followed by a back-migration from New Guinea to mainland East Asia. This putative back-migration would then have spread both New Guinea and Denisovan ancestry throughout East Asia and Siberia, and ultimately to the Americas. To distinguish between these two scenarios, we repeated the previous analysis but substituted Australians for New Guineans, comparing the archaic admixture in EE/NA populations (as a fraction of that in Australians) to the statistic *f*_4_(*Yoruba, Australian; French, X*). The results are virtually identical to those obtained with New Guineans as the comparison (Fig. 5*C*, 5*D*). Moreover, whereas Oceanian populations are more closely related to New Guineans than to Australians, EE/NA populations are equally related to Australians and New Guineans (Table S5). These results indicate that the archaic ancestry in EE/NA populations is shared with the common ancestor of Australians and New Guineans, and hence reflects the same admixture event.

## Discussion

Our analyses demonstrate that, in addition to being prevalent in Oceanian populations, Denisovan introgression is present in East Eurasian and Native American populations, even though the amount of Denisovan alleles in these latter populations is relatively small. These results thus confirm and extend previous studies suggesting Denisovan ancestry outside of Oceania (4-6). In particular, as found previously (2), Denisovan ancestry in Oceania is highly correlated with New Guinea ancestry. This suggests that these populations have either shared ancestry or contact with New Guinea that is more recent than the Denisovan admixture event. However, previous studies did not exclude the possibility that more ancient shared ancestry with New Guinean (after the Denisovan admixture event but before the divergence between New Guinean and Australian) explains the correlated signals of Denisovan and New Guinean ancestry in Oceania. Another potential explanation would be migrations from Australian rather than New Guinean, which could still produce a significant correlation between Denisovan and New Guinean ancestry as a consequence of the genetic relationship of Australians and New Guineans. To test these other possibilities, we compared amounts of Denisovan and Australian ancestry in Oceanian populations, and found that New Guinean ancestry does indeed provide a better explanation for the Denisovan ancestry in these Oceanian populations than does Australian ancestry (Table S5).

Our results also show a consistent signal of a low-level of Denisovan ancestry outside of Oceania, in populations of East Eurasia and the Americas. Although this signal does not reach significance in all populations (Fig. 4), given how widespread the signal is, it seems most reasonable to assume that all EE/NA populations probably do harbor some Denisovan ancestry. As with Oceanian populations, the Denisovan ancestry in EE/NA populations is correlated with their New Guinean ancestry. However, unlike Oceanian populations, the Denisovan ancestry in EE/NA populations is equally correlated with their Australian ancestry, and moreover EE/NA populations are just as closely related to Australians as they are to New Guineans (Fig. 5).

There are (at least) two potential scenarios that could account for these results. First, there was introgression from a population related to Denisovans into a modern human population that was ancestral to all EE/NA and Oceanian populations. After the separation of the ancestral EE/NA and Oceanian populations, subsequent migration(s) then brought other modern human ancestry into the ancestors of EE/NA (but not Australian or New Guinean) populations, thereby “diluting” Denisovan ancestry in EE/NA populations. This scenario has two important consequences. First, it means that the introgression between Denisovans and modern humans did not necessarily occur in island Southeast Asia as postulated previously (2), but instead could have occurred closer to the vicinity of Denisova Cave, in southern Siberia. Second, identifying the source(s) of the other modern human ancestry in EE/NA populations would be of great interest for further understanding the genetic history of human populations.

The second scenario that could explain the different amounts of Denisovan ancestry in Oceanians vs. EE/NA populations would be that Denisovan introgression occurred specifically in a population ancestral to Australians, New Guineans, and the Mamanwa, as hypothesized previously (2). After the Denisovan admixture, but before the divergence of Australians and New Guineans, there would then have been a back-migration from Oceania to mainland East Asia, which would have contributed both Denisovan and shared Australian/New Guinea ancestry to the ancestors of present day EE/NA populations. While we are not aware of any previous suggestions of such a back-migration from archaeological, anthropological, or genetic evidence, we also are not aware of any evidence that would disprove such a back-migration.

It thus seems that at present these two scenarios are equally plausible explanations for our results. In any event, the inescapable conclusion is that Denisovan ancestry is more widespread in modern human populations than thought previously to be the case, and moreover human genetic history must consequently also be more complicated than previously believed. Mapping the segments of Denisovan ancestry in modern human populations, as has been done for Neanderthal ancestry (5,16,17), should provide more insights into the history and consequences of the interactions between Denisovans and modern humans.

## Materials and Methods

We genotyped 168 individuals from 20 populations of Oceania and Southeast Asia with the Affymetrix Human Origins Array (8). An additional 2890 samples from 236 worldwide modern human populations, and high coverage sequences of two archaic hominins (Altai Neanderthal, 52× (5) and Denisovan, 31× (3)) were merged into the dataset. Principal component analysis was performed with EIGENSOFT (18) version 5.0.1. We performed PCA on chimpanzee and archaic hominin data, and then projected modern humans. Formal tests for archaic admixture in modern humans was performed by *f*_4_ statistics (8, 10). Admixture proportions were estimated by an *f*_4_*ratio* (8, 11). More details of the materials and methods can be found in supporting information.

## Acknowledgements

We thank David Reich for assistance with the production of the genotype data, and Kay Prüfer, Janet Kelso, and David Reich for helpful discussion and commenting on the manuscript. This research was supported by the Max Planck Society.

## References

1. Reich D et al. (2010) Genetic history of an archaic hominin group from Denisova Cave in Siberia. Nature 468:1053–60.

2. Reich D et al. (2011) Denisova admixture and the first modern human dispersals into Southeast Asia and Oceania. Am J Hum Genet 89:516–28.

3. Meyer M et al. (2012) A high-coverage genome sequence from an archaic Denisovan individual. Science 338:222–6.

4. Skoglund P, Jakobsson M (2011) Archaic human ancestry in East Asia. Proc Natl Acad Sci U S A 108:18301–6.

5. Prüfer K et al. (2014) The complete genome sequence of a Neanderthal from the Altai Mountains. Nature 505:43–9.

6. Huerta-Sánchez E et al. (2014) Altitude adaptation in Tibetans caused by introgression of Denisovan-like DNA. Nature 512:194–197.

7. Wall JD et al. (2013) Higher levels of neanderthal ancestry in East Asians than in Europeans. Genetics 194:199–209.

8. Patterson N et al. (2012) Ancient admixture in human history. Genetics 192:1065–93.

9. Price AL et al. (2006) Principal components analysis corrects for stratification in genome-wide association studies. Nat Genet 38:904–9.

10. Reich D, Thangaraj K, Patterson N, Price AL, Singh L (2009) Reconstructing Indian population history. Nature 461:489–494.

11. MoorJani P, Patterson N, Hirschhorn J (2011) The history of African gene flow into Southern Europeans, Levantines, and Jews. PLoS Genet 7, doi: 10.1371/journal.pgen.1001373.

12. Delfin F et al. (2012) Bridging near and remote Oceania: mtDNA and NRY variation in the Solomon Islands. Mol Biol Evol 29:545–64.

13. Duggan AT et al. (2014) Maternal history of Oceania from complete mtDNA genomes: contrasting ancient diversity with recent homogenization due to the Austronesian expansion. Am J Hum Genet 94:721–33.

14. Wollstein A et al. (2010) Demographic history of Oceania inferred from genome-wide data. Curr Biol 20:1983–92.

15. Duggan AT, Stoneking M (2014) Recent developments in the genetic history of East Asia and Oceania. Curr Opin Genet Dev 29C:9–14.

16. Sankararaman S et al. (2014) The genomic landscape of Neanderthal ancestry in present-day humans. Nature 507:354–7.

17. Vernot B, Akey JM (2014) Resurrecting surviving Neandertal lineages from modern human genomes. Science 343:1017–21.

18. Patterson N, Price AL, Reich D (2006) Population structure and eigenanalysis. PLoS Genet 2:e190, doi:10.1371/journal.pgen.0020190.

